# Mechanics of biomimetic free-standing lipid membranes: Insights on lipid chemistry and bilayer elasticity

**DOI:** 10.1101/2023.08.21.554126

**Authors:** Alessandra Griffo, Carola Sparn, Fabio Lolicato, Friederike Nolle, Navid Khangholi, Ralf Seemann, Jean-Baptiste Fleury, Martin Brinkmann, Walter Nickel, Hendrik Hähl

## Abstract

The creation of free-standing lipid membranes has been so far of remarkable interest to investigate processes occurring in the cell membrane since its unsupported part enables studies in which it is important to maintain cell-like physicochemical properties of the lipid bilayer, that nonetheless depend on its molecular composition. In this study, we prepare pore-spanning membranes that mimic the composition of plasma membranes and perform force spectroscopy indentation measurements to unravel mechanistic insights depending on lipid composition. We show that this approach is highly effective for studying the mechanical properties of such membranes. Furthermore, we identify a direct influence of cholesterol and sphingomyelin on the elasticity of the bilayer and adhesion between the two leaflets. Eventually, we explore the possibilities of imaging in the unsupported membrane regions. For this purpose, we investigate the adsorption and movement of a peripheral protein, the fibroblast growth factor 2, on the complex membrane.

## 1.0. Introduction

Lipid membranes form the cell membrane of almost all living organisms, such as eukaryotic cells, some prokaryotes, and several viruses. The membrane’s ability to separate intracellular and extracellular ionic media makes it a functional capacitor that confines cells and organelles and regulates multiple translocation mechanisms^1^. Together with the capacitive function, a set of physical properties of lipid membranes, including elasticity, tension, and curvature^2^, strongly affects many of the processes that are involved across the membrane. Examples are endocytosis, exocytosis, or conformational reshaping of proteins embedded in membranes^3^. Also phenomena occurring at the protein-lipid interface as protein binding^4^, oligomerization and insertion depend on the physical properties of lipid membranes and on their dynamic nature, which comprises processes, as e.g., curvature fluctuations and diffusion. Supported lipid bilayers (SLBs) have been, so far, for their stability, a versatile and demanded setup to easily assess a wide range of information^5^. The presence of the supporting substrate sacrifices, however, their dynamic nature and thus alters the reliability of observed processes occurring in and across the membrane^6,7^. Liposomes, as an alternative system to SLBs, allow to explore bilayer properties without disturbing these dynamic processes, yet suffer from limited experimental access^8^.

Moreover, membrane properties and functions depend also on chemical composition and polydispersity of the lipids present in the cell membrane^9–12^. Biomimetic membranes should therefore be heterogeneous and polydisperse in lipid composition and resembling the mimicked living systems^2^ as much as possible. Each lipid contributes to complex regulation mechanisms across the cell. Charged lipids, such as phosphatidylserine (PS) and Phosphatidylinositol 4,5-Bisphosphate PI(4,5)P_2_, can regulate the binding affinity of proteins by electrostatic interactions ^13,14^. Sterols and sphingolipids are in particular known to influence the arrangement of lipids^12,15^ affecting both mechanical and chemical properties of lipid membranes and the resulting interactions with the surrounding environment. Also, cholesterol (CL) has been shown to significantly affect the bilayer stiffness. In addition, CL laterally segregates in domains in the outer leaflet of the membrane together with sphingomyelin, causing a considerable impact on the mechanics of the bilayer. Sphingomyelin (SM) has, due to its features, such as a high degree of saturation, asymmetric molecular structure, and extensive hydrogen-bonding properties, a very important role as a structural parameter in biological membranes. Furthermore, due to its tendency to interact with sterols, sphingomyelin largely regulates the cholesterol distribution within cellular membranes. More importantly, the tendency to form strong hydrogen bonds between sphingosine backbones of adjacent SM molecules is associated with an increase in compactness affecting the bilayer rigidiy^16,17^.

In the current work, we prepared biomimetic membranes on substrates about 1 µm diameter circular holes. The pore-spanning membrane parts are a system between liposomes and SLBs featuring a better nanoscale observation than liposomes and a higher similarity to the natural situation than SLBs due to the absence of the solid support. These bilayer-model systems show water compartmentalization and bending rigidity similarly to liposomes^18^, but are still accessible to atomic force microscopy (AFM) investigation, allowing to study a wider range of phenomena, e.g., monitoring the kinetics of protein activity on top of membranes^9,19 20^. In order to design a system close to the cell membrane situation, we used a complex plasma membrane-like composition (PMC) of PC:PE:PS:PI:CL:SM:PI(4,5)P_2_ with ratios 33:10:5:5:30:15:2 for this study. Additionally, lipids from natural origin with a higher distribution of the fatty acid chain lengths compared to otherwise mostly employed synthetic ones were chosen for our biomimetic membranes (**Figure 1**).

**Figure 1.**
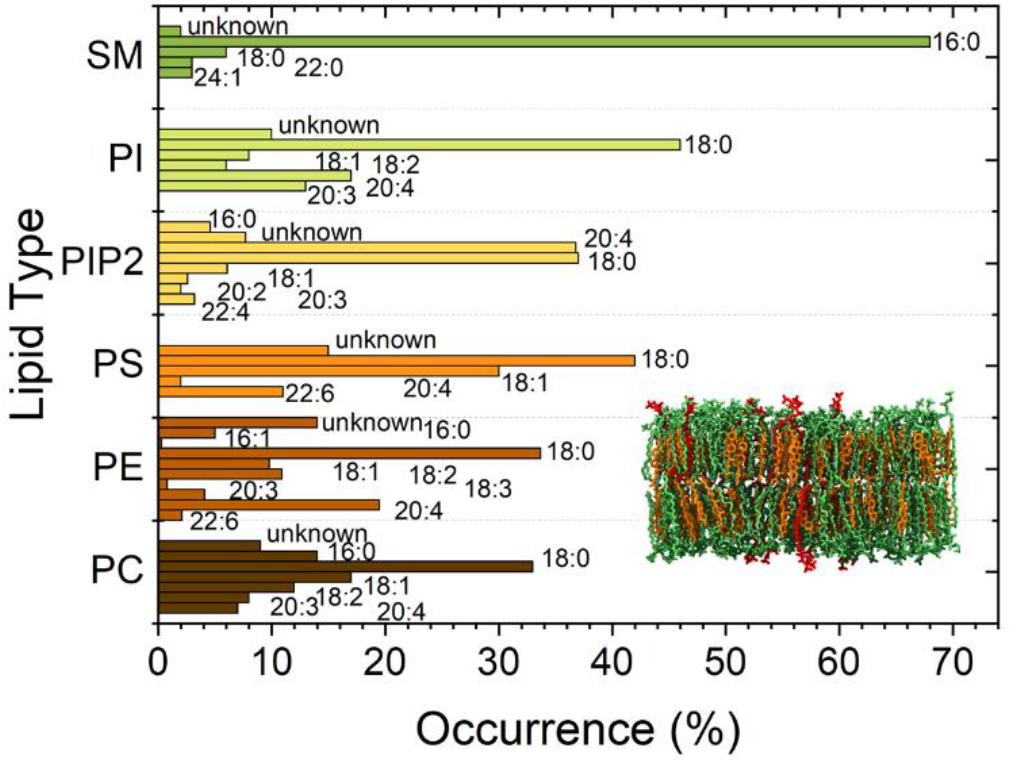
Fatty acid distribution of each of the natural lipids employed in the current study. The bars marked with X:Y show for a certain lipid type the fraction with chain length (number of C atoms) of the fatty acids X and number of unsaturated bonds Y. Lipid types are sphingomyelin (SM), phosphatidylinositol (PI), phosphatidylinositol bisphosphate (PIP_2_), phosphatidylserine (PS), phosphatidylethanolamine (PE), and phosphatidylcholine (PC). Data are adapted from Avanti Polar.

We investigated the effect of cholesterol and sphingomyelin on the membrane mechanics with the methods described below. In detail, we determined bilayer pre-stress and stretching elasticity for these heterogeneous membranes via AFM indentation and by analysis of optical micrographs of membrane standing in a microfluidic channel, while tuning the cholesterol and sphingomyelin content.

## 2.0. Materials and Methods

### 2.1 Preparation of Giant Unilamellar Vesicles

Unsupported membranes have been produced by bursting giant unilamellar vesicles (GUVs) on top of mercapto-1-undecanol functionalized gold TEM grids (Plano GmbH, pore diameter of 0.8 and 1.2 µm), by adapting the methods described in the literature (**Figure 2A**)^21–24^. Gold-coated TEM grids were functionalized with thiol self-assembled monolayers (SAMs). For that, they were first cleaned and made more hydrophilic via UV-Ozone treatment and then left overnight in a 1 mM solution of mercapto-1-undecanol in ethanol. After that, they were rinsed with ethanol to remove the unbound molecules and let dry. The functionalization was tested by water contact angle (WCA) measurements.

**Figure 2.**
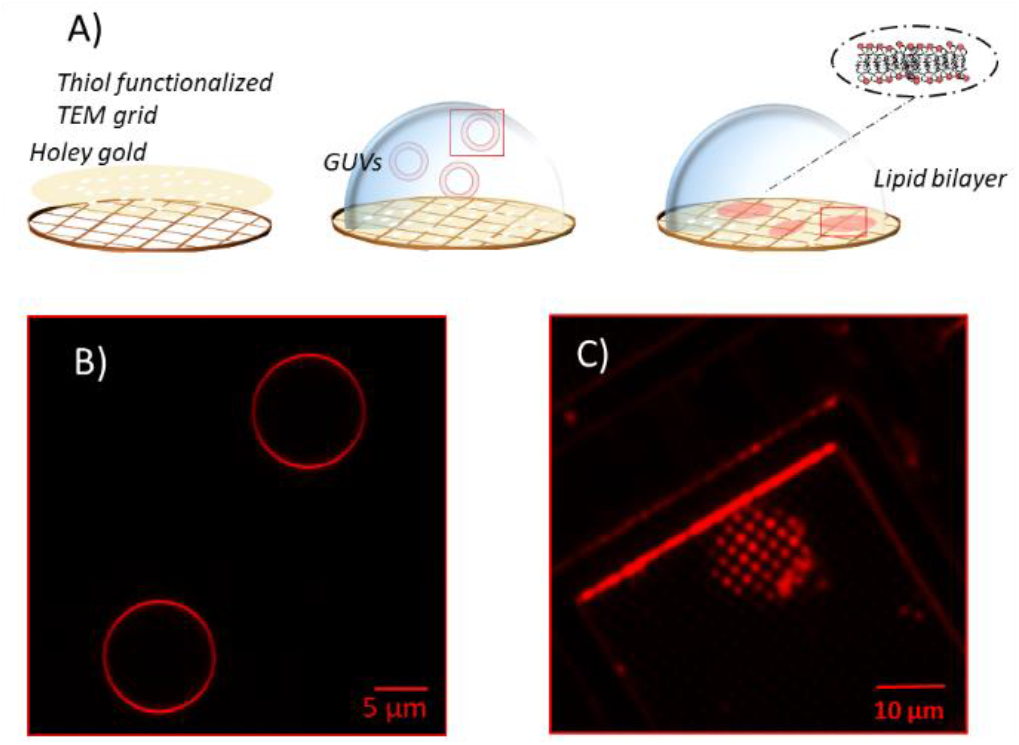
A) Sketch of the process of GUVs bursting on top of hydrophilic functionalized TEM grids and corresponding fluorescence microscope images of GUVs B) before and C) after bursting on TEM grid. Lipids are stained with Rhod-PE.

For creating the GUVs, a plasma membrane-like lipid composition (PMC) consisting of 30 mol% cholesterol (CL), 15 mol% sphingomyelin (SM), 33 mol% phosphatidylcholine (PC), 10 mol% phosphatidylethanolamine (PE), 5 mol% phosphatidylserine (PS), 5 mol% phosphatidylinositol (PI) and 2 mol % PI(4,5)P_2_ (Avanti Polar Lipids, Alabaster, AL) and a PMC lacking of CL and SF (PMC-) were prepared in chloroform with 1.5 mM concentration. All membrane lipids were purchased from Avanti Polar Lipids. They have been purified from natural extracts [bovine liver (PC, PE, PI), egg chicken (SM), porcine brain (PS, PI(4,5)P_2_), and ovine wool (CL)] and are therefore not uniform in alkyl chain length and unsaturated bond content (see **Figure 1**). The GUVs were generated by electroformation, according to the protocol of Steringer et al ^22^. In detail, 5 µL of the lipids dissolved in chloroform solution was casted on platinum (Pt) wires, dried and immersed in sucrose solution (0.3 M). An AC-voltage of 1.5 V and a frequency of 10 Hz were applied for 40 min. After that, the frequency was reduced to 2 Hz for 20 minutes to allow the detachment of the GUVs from the Pt wires. The vesicles produced in this way (**Figure 2B**) were collected, stored at 4 ºC, and used no longer than two days after production. Unsupported membranes were formed by diluting the stored GUV suspension in a ratio 1:50 in HEPES buffer.

### 2.2 Unsupported Membranes for AFM

Unsupported membranes were formed by diluting the GUVs suspension in HEPES buffer 20 mM, 300 mM NaCl, 25 mM MgCl2 and letting them burst, due to the difference in osmolarity between the sucrose inside the vesicle and the outer buffer, on the top of hydrophilic TEM grids mounted on a petri dish. Bursting of GUVs was checked by fluorescence microscopy in the presence of Rhod-PE, as reported in **Figure 2**. Eventually, for the AFM imaging and indentation experiments the solution was exchanged with imaging buffer [HEPES buffer 20 mM, 100 mM NaCl, 1 mM Ethylenediaminetetraacetic acid (EDTA)].

### 2.3 AFM imaging, force spectroscopy and data analysis

Atomic Force Microscopy (AFM) images and indentation experiments were acquired using a Dimension FastScanBio AFM (Bruker-Nano, Santa Barbara, CA, USA). MSNL-D probes (spring constant 0.03 N/m, resonance frequency 15 kHz, nominal tip radius 2 nm) were used for indentation experiments, while MSNL-D, SNL-D and PEAKFORCE-HIRS-F-B cantilevers (spring constant 0.03 N/m, 0.06 N/m and 0.12 N/m and resonance frequency 15 kHz, 18 kHz and 100 kHz, respectively, all probes purchased from Bruker AFM Probes, USA) were used for imaging. For each cantilever, the spring constant was calibrated by the thermal tune method. Peak Force Tapping® is used for imaging. During imaging, scan sizes varied from 500 nm to 20 µm. In the force curve measurements, the approaching and retracting curves were recorded with zero surface delay and a ramp size of 100–200 nm. For each indentation experiment, hundreds of force distance curves, (*F*-δ), were recorded on membrane standing holes; for these measurements, the homogenously coated holes were selected. All experiments were performed at room temperature (controlled at ca. 20 °C).

The data obtained from AFM indentation experiments were processed using custom-made MATLAB scripts for baseline correction and contact point determination, and the breakthrough force of the first rupture event was extracted. Pre-stress and elasticity of the membrane were determined by fitting a model to the data that was derived by Jin et al.^25^. The model is an analytical exact solution of the Föppl-von Karman approximation for the ‘circular clamped elastic membrane under point load with pre-stress’. It can thus describe situations with any pre-stress value within the two limiting cases with negligible pre-stress, which is characterized by a cubic dependency of the indentation force *F* on the indentation depth δ, and very large pre-stress, where the force is a linear function of the indentation depth. It is, however, not possible to write the general (*F*-δ) dependency in explicit form. Fitting the model to the data was therefore done using a custom-made MATLAB script (see SI for details).

### 2.4 Free-standing membranes in microfluidic channel

We employed an H-junction geometry previously tested as a platform to produce stable membranes to form unsupported lipid membranes in microfluidics ^26,27^, cf. **Figure 3A** and **B**. To guarantee a good wettability for the continuous oily phase in the microfluidic device, PDMS is used as material. The microfluidic devices are fabricated from a silicon elastomer base and curing agent (PDMS, Sylgard 184, Dow Corning, USA) mixed in a ratio 10:1 and are casted on a silicon wafer, having structures produced by photolithography with a SU-8 photoresist, and cured for 6 h at 65 °C. After curing the PDMS elastomer, it is peeled off from the mold and exposed to nitrogen plasma (Diener electronic GmbH, Germany) together with an also PDMS coated glass slide. The treated PDMS device is sealed with the glass slide and heated at 125 °C for 1–2 h to restore the hydrophobic properties of the PDMS. At the start of the experiment, this microfluidic device is filled with a continuous phase of squalene containing the lipids (5 mg/mL). Subsequently, two fingers of aqueous imaging buffer solution are injected into both channels of the device. Due to the wettability conditions, a thin squalene film remains between the buffer and the device walls, and the lipids contained in the squalene solution will decorate the oil-buffer interface forming a monolayer. A slow flow is created by hydrostatic pressure such that the lipid decorated oil-buffer interfaces are pushed through both channels and are brought in close vicinity at the junction. When the two monolayers touch each other, a zipping phenomenon driven by the intermolecular forces between the hydrophobic tails will occur, forming a bilayer (**Figure 3A, C-D**). Optical images of the bilayer formation occurring in the chip were acquired with Fluorescence Microscopy at 10x magnification. The angle θ between the two leaflets of the bilayer can be obtained by analyzing the optical micrographs.

**Figure 3.**
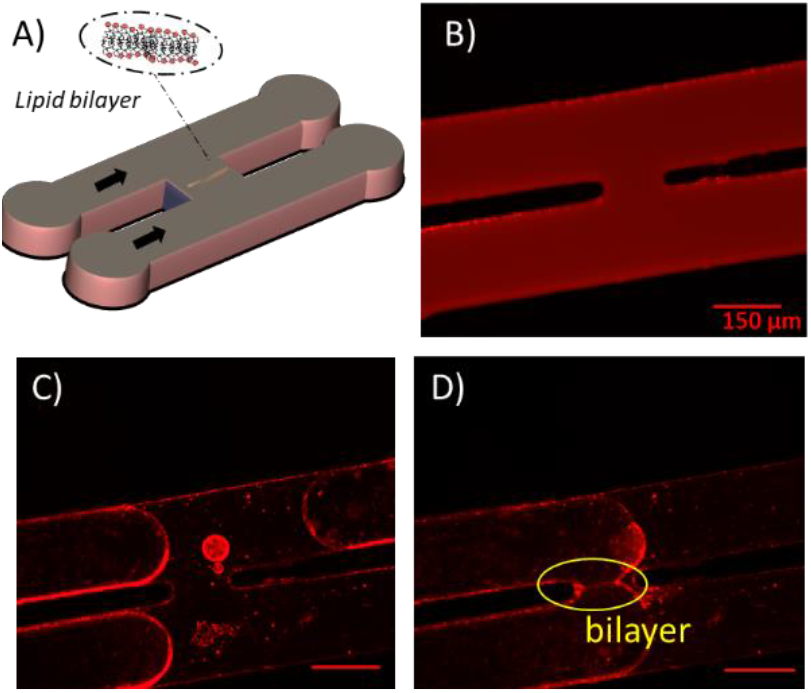
A) Sketch of the design of the microfluidic setup featuring a H-junction. B–D) Fluorescence microscope images (Lipids are stained with Rhod-PE) monitoring the process of lipid bilayer formation from (B) initial flooding of the device with lipid-squalene solution, (C) slow injection of the buffer phase and (D) contacting the oil-buffer interfaces coated now with a lipid monolayer forming a bilayer at the contact.

### 2.5 Interfacial tension γ and bilayer Tension Γ

Interfacial tension γ of a water/oil interface decorated with a monolayer of the lipid mixture is measured via the standard pendant drop method using a WCA goniometer (OCA 20, DataPhysics Instruments GmbH, Filderstadt, Germany). Briefly, a 1 mg/mL solution of lipid in squalene (0.858 g/cm^3^) is transferred in an optical glass cuvette. A drop of imaging buffer solution (ca. 1 g/cm^3^) is introduced from a needle that is dipped into the solution and the change of drop shape was observed over time, i.e. as a monolayer is formed at the interface. The droplet contour was fitted according to the Young-Laplace equation to obtain the interfacial tension γ. The decrease of γ due to the adsorption of lipids to the newly created interface is recorded over 20–30 minutes until a plateau is reached, denoting the interfacial tension of the monolayer decorated interface.

From the values of the interfacial tension γ and the bilayer contact angle θ, the bilayer tension Γcan be calculated using Young’s equation^28,29^ :

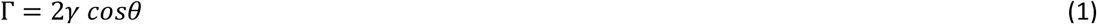

Although the bilayer tension Γ and the pre-stress in the spanned bilayer σ are, in principle, the same quantity, namely the in-plane stress in the bilayer, they are named differently here to clarify their different origin: as the preparation conditions are different, it is not expected to measure the same in-plane stress in both experimental setups. Particularly, the bilayer tension Γ in the microfluidic setup results from the interfacial tension γ and the adhesion energy ∆*W* between the two bilayer leaflets as

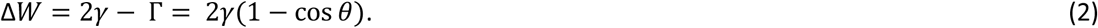

## 3.0. Results

Unsupported lipid membranes with PM-like composition (PMC) or with a PM-like composition without CL and SM molecules (PMC^−^) spanning hydrophilic coated TEM grids (pore diameter 1.2 µm and 0.8 µm) were formed. AFM scans (**Figure 4**) revealed the presence of membranes over the selected spots. The AFM cross sections (**Figure 4b** and **4c**) show that the membrane covers the rim around the hole, which has a height of ca. 20 nm, and is recessed inside the hole below the actual substrate’s surface to a depth of ca. 70–100 nm.

**Figure 4.**
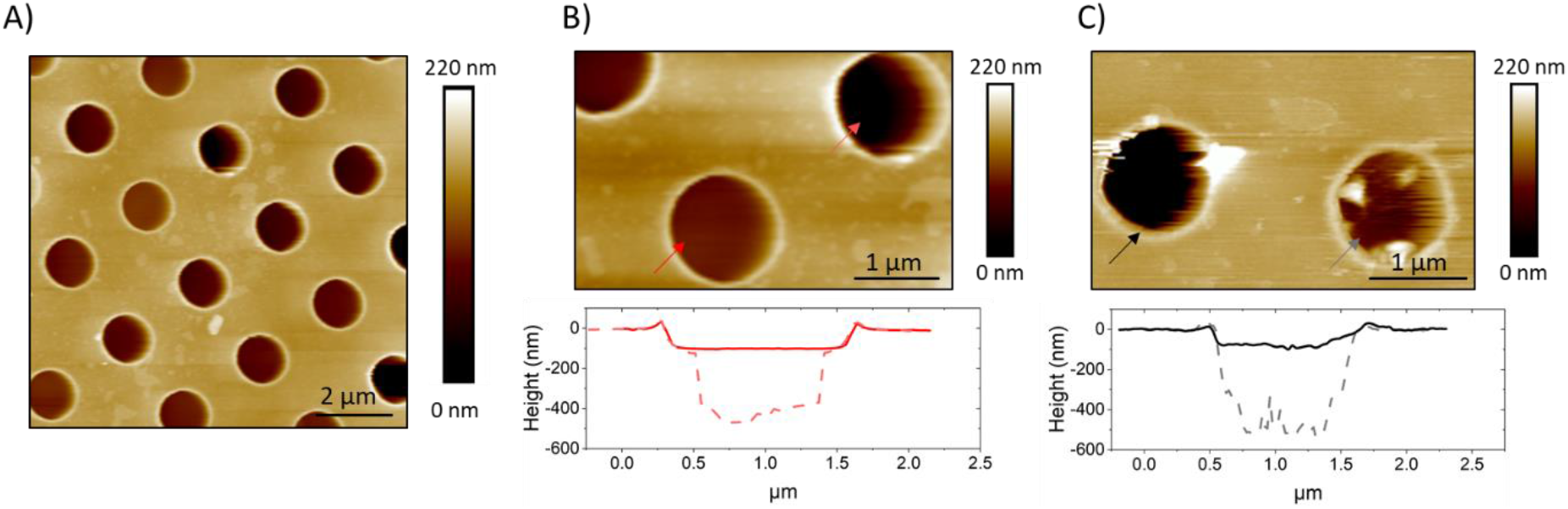
AFM images showing PMC (A, B) and PMC^−^ (C) lipid membranes spanning holes in a functionalized Au-grid. Section analysis in B and C show intact (red and black) and broken (light red and gray dashes) membranes respectively for PMC and PMC^−^. Images were recorded at a PeakForce Setpoint of 280 pN. Additional images of PMC and PMC^−^are reported in **Figure SI1**.

We performed AFM force-indentation (F-δ) measurements in the center of membrane-covered holes (**Figure 5A**). From these data, we extracted the three parameters of breakthrough force (BF), elastic modulus (E), and bilayer pre-stress (σ) in order to unraveled the effect of CL and SM on the mechanical properties of the PMC membranes. *BF* is defined as the maximum force the membrane can withstand before it ruptures, indicated by an abrupt decrease in force. It can be correlated to the packing density of lipids in the membrane^30^. The F-δ curves reported in **Figure 5A** show a nonlinear response in the region between the *snap in/contact point*, which is due to tip-surface interactions like capillary forces, and the *BF* point. This region was fitted (green dashed lines) with a model from non-linear elastic theory of membranes^25,25^ (**see SI**) enabling an estimation of both the elastic modulus *E* of the bilayer and its pre-stress σ.

**Figure 5.**
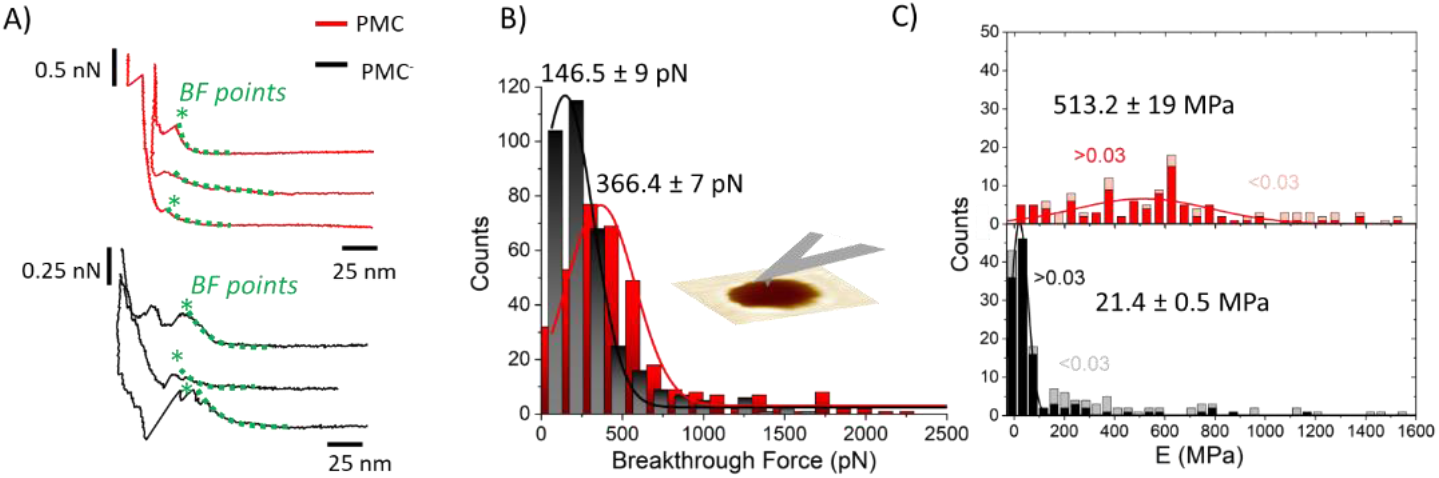
A) F-δ curves on PMC (red) and PMC^−^ (black) membranes. B) Histograms of the breakthrough force values (BF) for PMC (red) and PMC^−^composition (black) fitted using a Gaussian function. C) Stacked histograms reporting the Young’s modulus E for PMC (red squares for δ > 0.03 and pink for δ < 0.03) and PMC^-^ (black squares for δ > 0.03 and gray for δ < 0.03). Additional F-δ curves are reported in **Figure SI2**.

The histograms shown in **Figure 5B** are a collection of *BF*s from PMC and PMC^−^ membranes. They exhibit peaks that are fitted by Gaussian functions centered at (366 ± 7) pN for PMC and at (146 ± 9) pN for PMC^-^.

In **Figure 5C**, *E* values obtained from the fitting procedure are plotted. Since no dependency on the hole radius was observed, we combined data from membranes spanning holes of 0.8 µm and 1.2 µm diameter. The maximum indentation depth reached in the experiment affects, however, the accuracy of the estimation of the Young’s modulus. Since the stretching of the bilayer is more pronounced for larger indentations, the elastic modulus will be more important for the force response of the bilayer and thus, the estimation of *E* is improved. From a plot of the obtained *E*-values vs. the scaled maximum indentation depth on hole radius δ/R we decided for a threshold of δ/R = 0.03. Only values above this threshold were used for the estimation of the mean value of *E*. In **Figure 5C**, histrograms of the *E*-value distributions are presented where the disregarded values with δ/R < 0.03 are displayed as lighter shaded bars showing that they are mainly high value outliers in the distributions. Values above this threshold (red and black bars of **Figure 5C**) are fitted with a Gaussian distribution with a peak value for the elasticity modulus *E* at (513 ± 19) MPa and (21.4 ± 0.5) MPa for the PMC (red squares) and PMC^-^ (black squares) compositions, respectively.

Pre-stress values σ extracted from F-δ measurements that are shown in **Figure 5A** are reported in **Figure 6A**. We find that the value for σ is reduced from (5.2 ± 0.8) mN/m for the PMC membrane to (2.0 ± 0.2) mN/m for the PMC^−^ membrane. For qualitative comparison, the tension in the bilayer for both compositions was also determined from the contact angle θ of two contacted lipid monolayers in a microfluidic setup. Together with the interfacial tension γ of the monolayer, which was measured using the drop shape analysis of a pendant drop, and employing the Young Laplace equation (eq. 1)^26^, bilayer tension values Γ of (13 ± 4) mN/m and (10 ± 1) mN/m (**Figure 6B**) were obtained for PMC and PMC^−^ membranes, respectively. These values relate via eq. (2) to adhesion energies ∆W between the sheets of (2.3 ± 0.2) mJ/m^2^ and (1.4 ± 0.3) mJ/m^2^.

**Figure 6.**
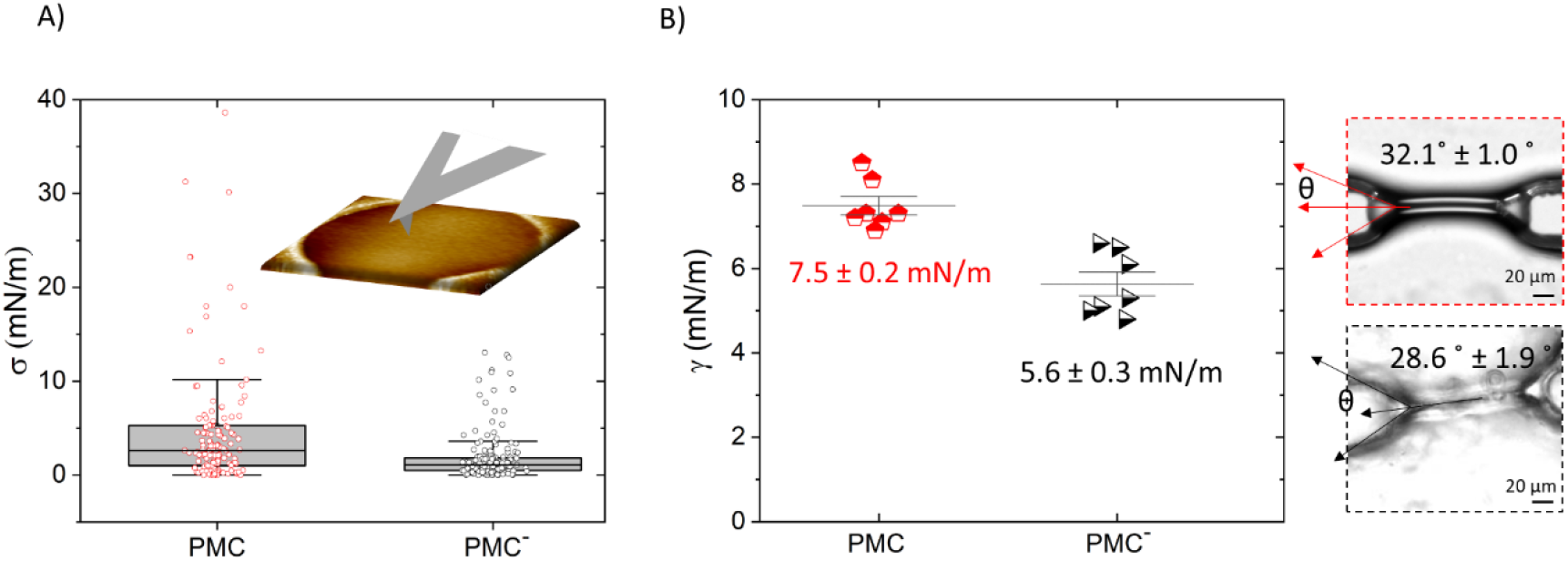
A) Box plots of pre-stress σ (± SE) values recorded via AFM indentation measurements for PMC and PMC^−^ compositions. B) plots of interfacial tensions recorded via pendant drop measurements and optical micrographs of the membranes formed in a microfluidic (µFlu) chip showing respectively the interfacial tension γ and the bilayer angle θ for PMC and PMC^-^ compositions. θ and γ are used to calculate the bilayer tension Γ with the Young Laplace equation.

In addition to the mechanistic insights, we used the imaging capabilities of the AFM to resolve the bilayer surface. Our choice to employ unsupported membranes for AFM investigation is guided by the need to construct platforms for protein observation and detection of its related processes, i.e., diffusion, kinetics assembly, pore forming mechanisms. **Figure 7** shows a series of AFM images of the free-standing part of the bilayer recorded while the protein fibroblast grow factor 2 (FGF2-GFP) was added to the solution above the bilayer, using a fluid cell setup. These AFM time-lapse frames show the adsorption of this peripheral protein across the bilayer. However, due to the fluidity of lipid membranes and the difficulty of imaging standing portions of lipid membranes, it is hard to picture a clear statement from the reported images.

**Figure 7.**
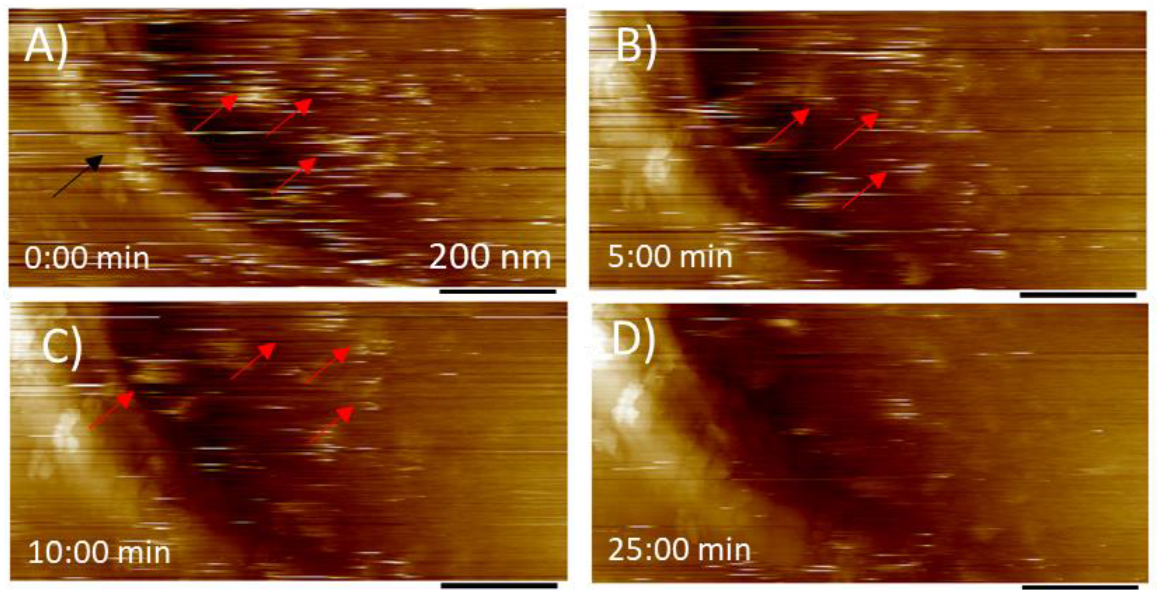
**A-D)** Height AFM time-lapse frames recorded under flow while FGF2-GFP proteins are injected. FGF2-GFP proteins are observed overtime on the free-standing portion of the membrane. Scale bar is 200 nm and z height 80 nm. The black arrow points to the rim of the membrane spanned hole and red arrows point some of the proteins.

## 4.0. Discussion

The described characterization allowed us to gain insight to the mechanics of membranes. The *BF* values of (366 ± 7) pN and at (146 ± 9) pN recorded by AFM for both unsupported PMC and PMC^−^ membranes imply a difference in the compactness of the two membranes. The PMC membranes exhibit a higher breakthrough force, meaning that the membrane can resist the applied force more strongly. This observation matches quite well with the condensing effect of cholesterol^31^, which is well known to regulate the membrane fluidity and produce an ordering of the lipids, leading to a decrease in its permeability^31–33^. Such insights have been suggested already by studies of SLBs^34–36^ where information about the chemical composition of the membrane, its lipid packing, and the surrounding environment were gained from the *BF*. Sullan et al.^37^ found that an increase of the *BF* was observed in the SM/Chol-enriched liquid-ordered domains (Lo) compared with the DOPC-enriched fluid-disordered phase (Ld). While the *BF* values they reported for SLBs were in the nN-range, our results for free-standing membranes fall, however, in the pN-range. Such differences are reasonably explained by the absence of a solid support. In our setup, the applied stress from the AFM cantilever causes a stretching of the membrane that, in turn, causes a reduction of the packing density until the inter-molecular distances are large enough to let the bilayer rupture at the *BF* point. In the SLB, the AFM tip has to push the molecules in order to punch through the bilayer, inducing an even higher packing density in the surrounding.

A closer insight into the mechanics of PMC lipid membranes and into the interplay of cholesterol and sphingomyelin can be obtained by extracting the stretching elasticity of the bilayer from the F-δ curves. Therefore, a theoretical description derived from the von the Karman Föppel model of elastic membranes was used. For a clamped elastic membrane under a central point load, a linear relation between force and indentation is predicted if the indentation is small and the pre-stress is not negligible. For higher indentations and small or negligible pre-stress, stretching of the membrane dominates and leads to a cubic relation depending on the elasticity for the membrane^38^, *F* ∝ *E* δ^3^. As an approximation, a linear superposition of those limiting behavior descriptions is used in similar studies^39–41^

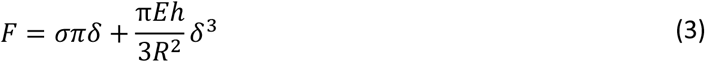

where *h* is the bilayer thickness, here assumed as 6.7 nm for PMC membranes and as 6.0 nm for PMC^−^ membranes^42,43^, *R* the hole radius, *E* the elastic modulus and *σ* the pre-strain of the membrane. Note, however, that in this setup the membrane is stretched laterally and thus only its lateral 2D elasticity *E*^*2D*^ is probed. *E*^*2D*^ is related to *E* as *E*^*2D*^ = *E* ⋅ *h* under the assumption of an isotropic material, which is not strictly valid in this case. We nevertheless state the values for *E* for a better comparability.

In the data of this study (**Figure SI3 and SI4**), both behaviors can be observed: a constant slope for low indentations and a deviation from the linear dependency for higher indentations of δ/*R* ≳ 0.03. Therefore, the accuracy with which *E* is determined increases with higher δ. Physically, this reflects the notion that the more the membrane is indented, the more it is stretched, and the more important Young’s modulus becomes.

Since the data of this study fall, however, mostly in the transition region between linear and cubic behavior, where the inaccuracy of the simple model is highest, we chose a slightly more complicated, yet analytically exact solution of the theoretical model by Jin et al.^25,44^ (see SI for details). Data shown in **Figure 5C** and **6A** were obtained by fitting this model to the experimental *F*-δ curves. Compared to the approximation of eq (1), we obtained slightly lower values for *E* and *σ* as well as a lower scattering (a comparison between the approximate and exact solution of the model is reported in **Figure SI3**). The obtained elasticity values recorded upon membrane indentation vary from 10 to 1000 MPa. Thereby, lower elasticity values are found for higher maximum indentation depths, with the smallest near 100 and 30 MPa for PMC and PMC^−^ membranes, respectively (**Figure SI3**). The latter values correspond to the maximum achievable membrane stretching.

The obtained results on membrane mechanics match with many works reported in literature^15,45^ where the role of SM and CL is attributed to an increase in stiffness. Yet, in most of the studies elasticity of membranes has been extensively probed for supported lipid bilayers and vesicles, but rarely for unsupported membranes. To enable a comparison with elasticity measurements from vesicle indentation^46,47^, where normally the bending rigidity is probed, we calculated this property from our measurements via

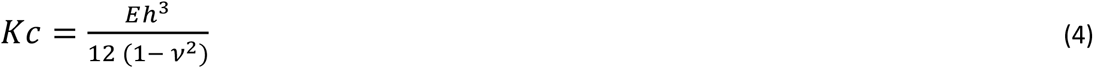

where υ is the Poisson ratio kept as 0.485^48^, *E* the elastic modulus and *h* the membrane thickness. We find values of about 200*10^−19^ J and 9*10^−19^J at maximum indentation depths for PMC and PMC^−^ membranes, respectively. Such values are slightly above those typically reported in the literature for membranes not containing rigid molecules such as SM and CL ^9,20,49^ and could be attributed to the presence of rigid-like lipids such as SM and CL that increase the membrane rigidity. Our results report a remarkable membrane rigidity, which is even larger compared to cholesterol enriched membranes ^15,50^. Sphingomyelin, in fact, increases the packing density of the membrane due to its hydrogen bond forming capacity with neighboring lipids^17^. Also, the PI(4,5)P_2_ may cause, due to its conical shape, a stiffening of the membrane^15^ and an increase of the lateral pressure in the acyl chain region^51^.

Studies on tension and pre-stress instead have been carried out more extensively also for monodisperse and unsaturated lipids on unsupported membranes^21,23,52^. In the work of Kocun et al.^21^, site specific force indentation experiments on porous substrates were performed to determine membrane tension as a function of lipid composition. Although they used a similar setup, their pure POPC or DOPC membranes showed only a linear force-indentation behavior, which did not allow for a determination of the bilayer’s elasticity. They obtained, however, an apparent spring constant, which depends on the preparation-imposed pre-tension of the bilayer and is comparable to (*σ*π) in our analysis. They observed an increase in the apparent spring constant for DOPC membranes from about 1.0 mN/m to 3.5 mN/m after the addition of cholesterol, which matches well with our findings, where the addition of the two rigid-like components cholesterol and sphingomyelin led to a shift from 2 mN/m to about 5 mN/m. In the case of the spanned membrane, the tension of the membrane is influenced by the adhesion to the substrate surrounding the hole, where a higher adhesion should lead to a higher tension^53^. For the membranes in the microfluidic setup, the tension is directly given by the adhesion between the sheets via eq. (2). For both situations, a higher tension thus means an increase in adhesion; in one case, to the substrate and, in the other case, between the leaflets. It is interesting to note that the addition of CL and SM in the membrane leads, in both cases, to an increased adhesion. However, the ratio of this increase seems too large to be attributed solely to an increased molecular density in the bilayer.

Our observations shed light on the function of sterols and sphingolipids on more complex compositions presenting larger lipid heterogeneity and fatty acid distribution as the degree of unsaturation, and polydispersity can tune the properties usually attributed to CL and SM^45^. Previous studies showed that for certain proteins, i.e. the FGF2 used in our experiments, both binding and translocation are highly enhanced when cholesterol and sphingomyelin are present^54^. The underlying mechanism remains not entirely elucidated, but it’s probable that variations in tension and the lipids’ inherent properties play significant roles in this phenomenon.

## 5.0. Conclusions

The technology of membrane biomechanics is continuously in evolution and currently many systems are studied, from in vitro constructed liposomes^55^, pore spanning membranes^35^, synthetic cells^56^ to entire cells^57^, using a variety of techniques, including AFM, fluctuation spectroscopy and micropipette aspiration. In our study, we employed Atomic Force Microscopy, Atomic Force Indentation and less extensively also microfluidics, to achieve insights into mechanical properties, such as bilayer tension and elasticity, morphological features, and protein-membrane interaction studies on unsupported membranes. Starting from the pore spanning technology we used biomimetic lipid membranes for which we chose a plasma membrane-like composition from natural lipids, to shed light on the properties of complex membranes, which feature lipid heterogeneity and polydispersity. The focus was mainly on the role of cholesterol and sphingomyelin, for which there is still a lack of information. The shift in the values of elastic modulus, of the breakthrough force as well as in the adhesion energies between PMC and PMC^−^ shows how the combination of different bilayer components affects the compactness and the mechanics of the membranes. This led us to the conclusion that cholesterol and sphingomyelin increase membrane tension and rigidity even in more complex PMC compositions. Eventually, we propose the described unsupported membranes as an alternative to supported ones as model for systematic studies with more natural dynamics.

## Author Contributions

A.G. performed the experiments and designed the work, F.N., N.K., J.-B. F. helped for the microfluidic experiments, C.S., F.L. helped in vesicle productions and W.N. supported the scientific discussion and in planning the project, M.B. helped in choosing the mathematical model, H.H. helped in the data analysis and designed the work. All the authors contributed to the manuscript writing. A.G. and H.H are shared corresponding authors.

## Acknowledgements

This study was supported by the German Research Foundation in the framework of the CRC1027 (B2, B4), DFG Ni 423/10-1; WN, DFG LO 2821/1-1; FL, and Saarland University within the funding program Open Access Publishing and was conducted within the Max Planck School Matter to Life supported by the German Federal Ministry of Education and Research (BMBF) in collaboration with the Max Planck Society.

