## Supplementary material for "Mechanics of biomimetic free-standing lipid membranes: Insights on lipid chemistry and bilayer elasticity"

**Free-standing membranes for AFM investigation**

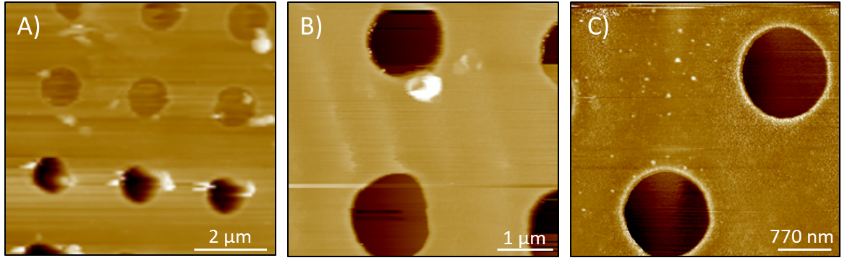

**Figure SI1.** Additional AFM height pictures for PMC^-^ (A, B) and PMC (C). Z heights are, 1 µm, 597 nm, 118nm respectively for A), B), C).

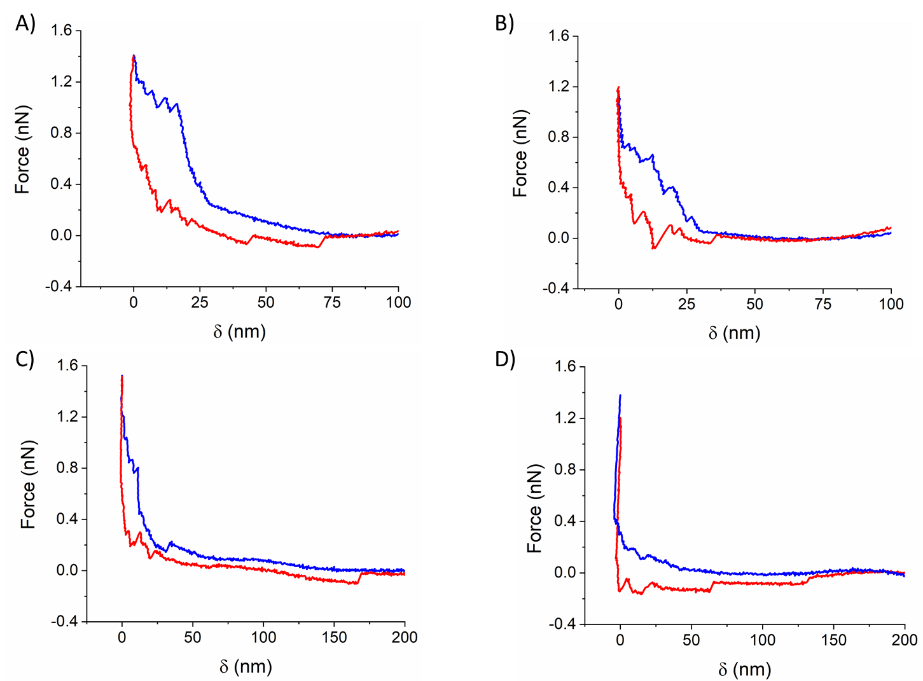

**Figure SI2.** Additional F-δ curves of lipid membranes having PMC A, B) and PMC^-^ composition C, D).

**Free-standing membranes for AFM investigation: analysis of FD curves**

The force-indentation curves were analyzed by a model derived from membrane theory following the solution of Jin et al.^1,2^. They derived an exact solution for a circular clamped membrane under central load from a circular or point-like indenter including residual stress. Their derivation is based on the non-linear governing equations for a Föppl-Hencky membrane, i.e. under the assumption that the stress-strain response of the material is still Hookean and rotations are sufficiently small. These equations are

|  | $r\frac{d}{dr}\left[ \frac{d}{r dr}(r^{2} N_{r}) \right]= -\frac{hE}{2}\left( \frac{dw}{dr} \right)^{2}$ | (S 1) |
| --- | --- | --- |
|  | $N_{r}\frac{dw}{dr}= -\frac{F}{2\pi r}$ | (S 2) |

, where $N_{r}(r)$ is the stress in radial direction, $r$ the radial coordinate, $w(r)$ the vertical displacement of the membrane at position $r$, $h$ the thickness of the membrane, $E$ the Young’s modulus of the membrane material and $F$ the applied load. The hole radius is $R$ and the indentation in the experiment corresponds to the displacement at the center $\delta:=w\left( 0 \right).$ The pre-stress σ is considered as the residual, radial stress when $F=0.$ It creates a radial displacement at the edge of the hole ($r=R$), which is used as prescribed parameter in the model and, together with $w=0$ at $r=R$, is the boundary condition for this system. The solution for this model, which Jin et al. derived (eq. (90) in Jin *et al.*^1^) consists of the two equations

|  | $\delta^{3}=F \frac{4R^{2}}{\pi Eh} \frac{\ln^{3} \left( \cot\frac{\theta_{m}}{2} \right)}{2\left( \frac{\cot\theta_{m}}{\sin\theta_{m}}-\ln\left( \cot\frac{\theta_{m}}{2} \right) \right)}$ | (S 3) |
| --- | --- | --- |
|  | $1-\nu=2\frac{\cos\theta_{m}-\sin^{2} \theta_{m} \ln\left( \cot\frac{\theta_{m}}{2} \right)-\cos^{3} \theta_{m}}{\sigma\left( \frac{16 R^{2}\pi^{2}}{Eh F^{2}} \right)^{\frac{1}{3}}\cos\theta_{m} \left( \cos\theta_{m}\sin\theta_{m}-\sin^{3} \theta_{m}\ln\left( \cot\frac{\theta_{m}}{2} \right) \right)^{\frac{2}{3}}-\cos^{3} \theta_{m}}$ | (S 4) |

, where $\theta_{m}$ is an integration constant with $0<\theta_{m}<\frac{\pi}{2}$ to be determined for every combination of $F$ and $\delta$. $\nu$ is the Poisson ratio, for which we used 0.485^3^ for this study.

For the ease of use and better numeric stability, we substituted $\beta=\cot\theta_{m}$ ($-\infty<\beta<\infty$) and rearranged the two equations to

|  | $F=\frac{\sigma^{\frac{3}{2}} 4\pi R}{\sqrt{Eh}} \left( 1-\nu\right)^{\frac{3}{2}} g_{1}\left( \beta\right)$ | (S 5) |
| --- | --- | --- |
|  | $\delta=\frac{\sqrt{\sigma} 2R}{\sqrt{Eh}} \left( 1-\nu\right)^{\frac{1}{2}} g_{2}\left( \beta\right)$ | (S 6) |

with

|  | $g_{1}\left( \beta\right)=\frac{\beta\sqrt{1+\beta^{2}}-\ln\left( \sqrt{1+\beta^{2}} +\beta\right)}{\left( 2-2\frac{\sqrt{1+\beta^{2}}}{\beta}\ln\left( \sqrt{1+\beta^{2}}+\beta\right)+\left( 1-\nu\right)\beta\right)^{\frac{3}{2}}}$ | (S 7) |
| --- | --- | --- |
|  | $g_{2}\left( \beta\right)=\frac{\ln\left( \sqrt{1+\beta^{2}} +\beta\right)}{\left( 2-2\frac{\sqrt{1+\beta^{2}}}{\beta}\ln\left( \sqrt{1+\beta^{2}}+\beta\right)+\left( 1-\nu\right)\beta\right)^{\frac{1}{2}}}$ | (S 8) |

Thus, for a given combination of the setup parameters $R$ and $h$ and the material properties $\sigma$, $E$, and $\nu$, the integration constant $\beta$ can be determined for any $\delta$ and the corresponding force $F$ calculated or vice versa. Implemented in a Matlab program, we used $\delta$ to calculate $\beta$ and hence $F$. While $R$, $h$ and $\nu$ were kept constant for a given experimental $F\left( \delta\right)$-curve, σ and $E$ are then adjusted in an iterative fitting process to obtain a best-fitting model curve.

Jin et al. derived also a solution for a circular indenter with radius c instead of a point-like indenter. This involves, however, determining a second integration constant and thus a more complicated solution and moreover renders the numerical fitting process more time-consuming. Based on some tests performed on a subset of the experimental curves, we could, however, not find significant differences in the obtained values for the fit parameters between the circular or point-like indenter model. With a tip radius of $r_{\mathrm{tip}}\approx5 \mathrm{nm}$, and thus $r_{\mathrm{tip}}\ll R$, the simplification seems therefore applicable, as expected.

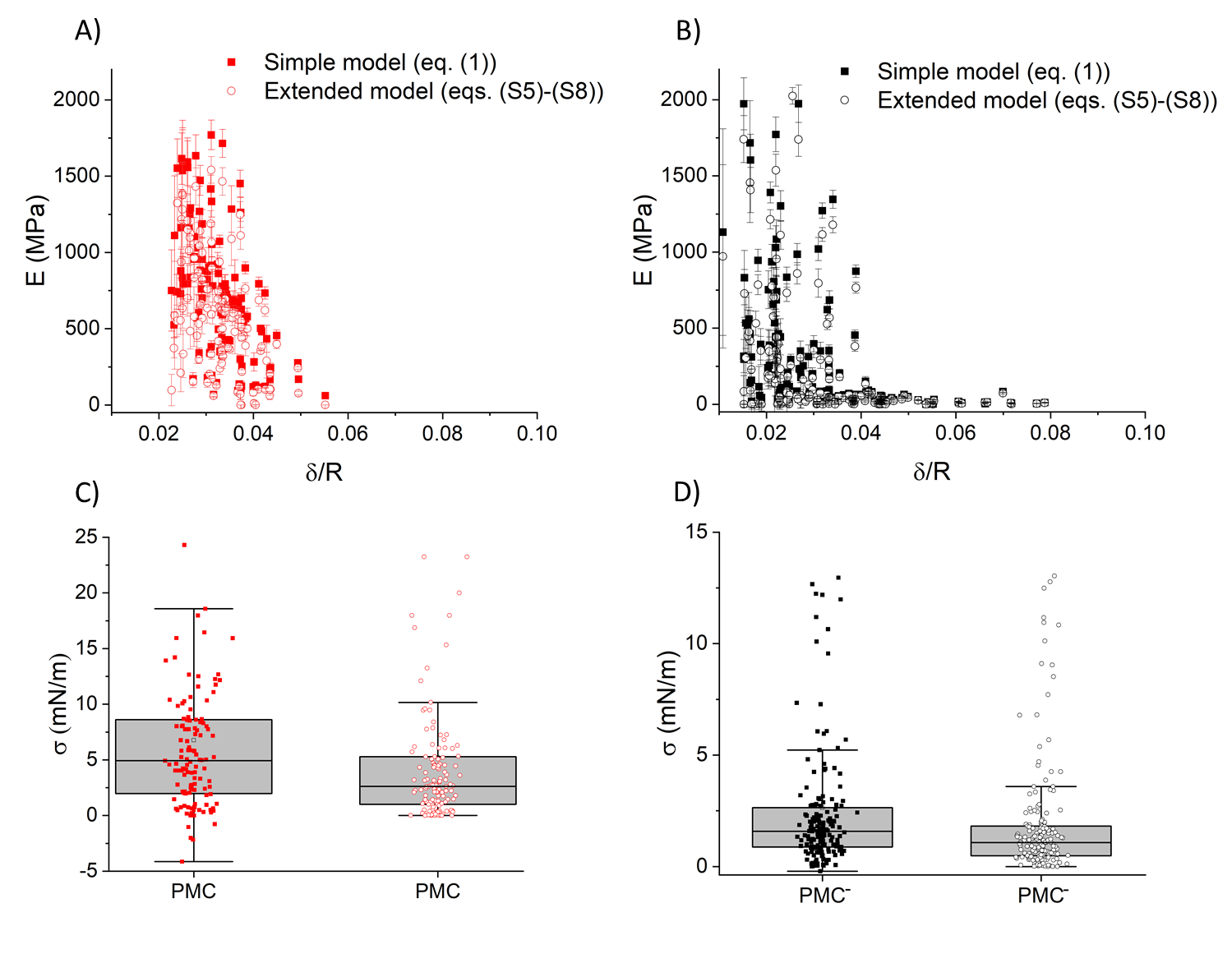

**Figure SI3**. (A, B) Elastic modulus *E* and (C, D) pre-stress *σ* as obtained from fitting the experimental force-distance curves. Filled square symbols are from fitting with the simpler model given as eq. (1), open circle symbols were obtained from fitting with the Jin et al. model (eqs. (S5)-(S8)). In (A) and (C) data from PMC membranes are shown, in (B) and (D) data from PMC^-^. (A) and (C) display a dependency of the obtained values on the normalized indentation depth as described in the main manuscript. The Data for $\delta/R<0.03$ are not used for mean value calculation. For (D) and (E) a consistent pre-stress *σ* dependency from $\delta/R$ is not observed. Mean values of pre-stress *σ* for PMC are 6.8 (filled squares) and 5.2 mN/m (open circles) while for PMC^-^ are 2.6 (filled squares) and 2.0 mN/m (open circles) for PMC^-^, showing a systematic shift toward lower E values when the Jin et al. model (eqs. (S5)-(S8)) is applied.

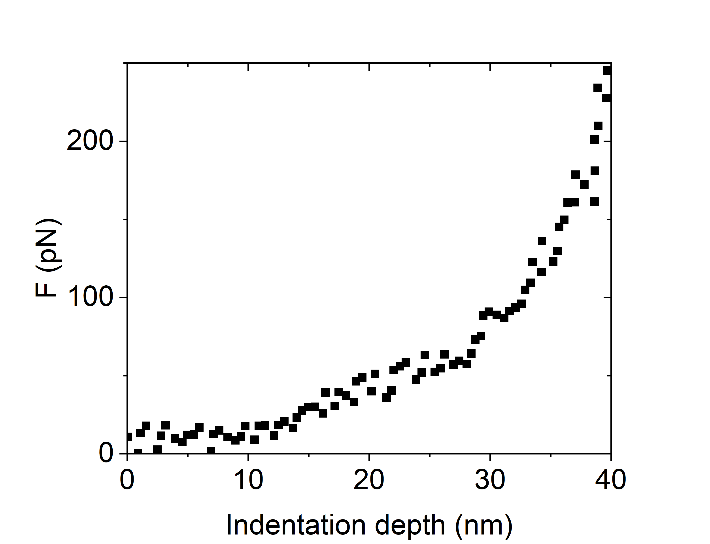

**Figure SI4.** F-δ curve for PMC^-^ showing a constant slope for low indentations and a deviation from the linear dependency for higher indentations of δ/R ≳0.03.
